# The Gulf Coast tick, *Amblyomma maculatum* (Ixodida: Ixodidae) and spotted fever group *Rickettsia* in the highly urbanized northeastern US

**DOI:** 10.1101/2021.12.08.471803

**Authors:** Waheed I. Bajwa, Leonid Tsynman, Andrea M. Egizi, Rafal Tokarz, Lauren P. Maestas, Dina M. Fonseca

## Abstract

We report the multi-year collection of the Gulf Coast tick, *Amblyomma maculatum* Koch (Acaridae: Ixodida: Ixodidae) in Staten Island, New York City (NYC) as well as their detection in Brooklyn, NYC, and in Atlantic and Cumberland counties in southern NJ, USA. The first detections on all sites were of adults but in Freshkills Park on Staten Island larvae were collected in a following year. Based on known observations on birds of this tick species, it is likely *A. maculatum* are expanding north on migratory birds, which are now often seen in Freshkills Park. The presence of larvae indicates that adults are being successful at finding hosts in Staten Island. We describe the landscape features of the area in Staten Island where populations were highest and larvae were detected, which could have facilitated the establishment of *A. maculatum*. Notably, we also report the presence of human pathogens *Rickettsia parkeri* in 5/10 (50%) of adults tested and *R. felis* in 1/24 (4.17%) of larvae tested. In addition to established populations in Staten Island we found evidence of *A. maculatum* in NJ and other NYC boroughs, suggesting current or future establishment is possible. The failure thus far to detect established populations in these areas may be due to inherent difficulties in detecting low density, spatially heterogeneous incipient populations, which could require targeted surveillance efforts for this species. We discuss the consequences to public health of the establishment of *A. maculatum* and detection of two additional rickettsial pathogens in the densely populated Northeastern US.

## Introduction

Ticks are disease vectors of increasing public health importance (Bouchard et al. 2019) responsible for the transmission of approximately 95% of vector-borne diseases currently circulating in the United States (Eisen et al. 2017). In the northeast US, most tick-borne diseases are transmitted by the blacklegged tick (*Ixodes scapularis* Say) although the lone star tick (*Amblyomma americanum* L.) is of increasing concern (Jordan and Egizi 2019, Molaei et al. 2019) and a new exotic species, the Asian longhorned tick (*Haemaphysalis longicornis* Neumann), first detected in New Jersey (NJ) in 2017, is now widely established (Beard et al. 2018, Rainey et al. 2018). Furthermore, two species native to the southeastern US, *Ixodes affinis* Neumann and *Amblyomma maculatum* Koch, appear to be expanding their distributions northward (Nadolny et al. 2011, Maestas 2013, Florin et al. 2014). Knowledge of which disease vectors are present is important to assess the risk to humans from the pathogens they are competent to carry.

*Amblyomma maculatum* (**Figure 1**) is an active, aggressive, generalist tick species (Nadolny and Gaff 2018). All life stages can feed on humans, but adults are most often encountered (Paddock and Goddard 2015). This species is a known vector of *Hepatozoon americanum*, the agent of American canine hepatozoonosis, a potentially debilitating disease of canines (Little et al. 2009), and is the primary vector of *Rickettsia parkeri*, the agent of a human disease variously called *R. parkeri* rickettsiosis, American Boutonneuse Fever or Tidewater Spotted Fever (Paddock and Goddard 2015). Although *Rickettsia parkeri* rickettsiosis is less severe to humans than Rocky Mountain spotted fever (caused by congener *Rickettsia rickettsii*) it nonetheless often requires antibiotic treatment (Paddock and Goddard 2015, Eisen et al. 2017, Silva-Ramos et al. 2021) and infected bites leave a characteristic eschar, i.e., a necrotic lesion at the site of the bite (Drexler et al. 2020). While only 39 confirmed cases of *R. parkeri* rickettsiosis had been reported in the US as of 2016 (Straily et al. 2016) it has been estimated that as much as a third of presumptive ‘Rocky Mountain spotted fever’ cases reported in the eastern US may actually be attributable to *R. parkeri*, due to lack of specificity of diagnostic tests (Raoult and Paddock 2005, Raoult and Parola 2008). Although *R. parkeri* may circulate in some small rodent species (Cumbie et al. 2020), very high rates of transovarial transmission (average of 83.7% in one study) appear to implicate *A. maculatum* as the primary reservoir of *R. parkeri* and allow the pathogen to persist in areas lacking competent vertebrate hosts (Wright et al. 2015). Other *Rickettsia* spp. have been detected in *A. maculatum* although its role as a reservoir or vector for these species is not clear. Candidatus *Rickettsia andeanae*, a species of undetermined pathogenicity, has been detected in *A. maculatum* from the central Midwest and Southeast US (Fornadel et al. 2011, Jiang et al. 2012, Paddock et al. 2015, Allerdice et al. 2019) often at very high prevalence. One study found that 73% of *A. maculatum* from Oklahoma were infected with Candidatus *R. andeanae* but none with *R. parkeri*, leading to a hypothesis that the presence of *R. andeanae* in a tick population may exclude *R. parkeri* (Paddock et al. 2015). *Rickettsia felis*, a human pathogen transmitted primarily by cat fleas (*Ctenocephalides felis* Bouché) has also been detected in a wide range of arthropods around the world (Pérez-Osorio et al. 2008) including two *A. maculatum* from the US (Jiang et al. 2012).

**Figure 1.**
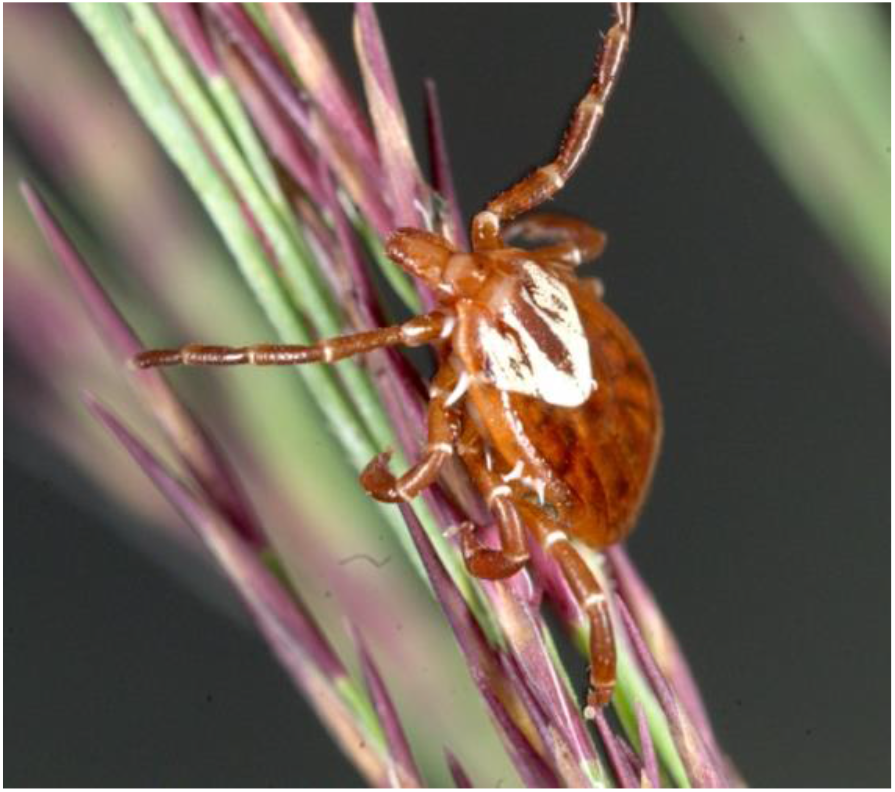
Female *A. maculatum* collected from *Phragmites* in Cumberland Co. NJ.

While *A. maculatum* was originally limited to the southeastern United States (Teel et al. 2010), it had been documented in Delaware in 2013 (Florin et al. 2014) where it was recently found established statewide (Maestas et al. 2020). There have also been recent detections of established populations in coastal Connecticut (Molaei et al. 2021) and in Illinois (Phillips et al. 2020). Migratory songbirds (passerines) have been implicated in the dispersal of immature (larvae and nymphs) *A. maculatum* during migration (Scott and Durden 2015) and it is thought that this association may be an important driver of its northward expansion (Paddock and Goddard 2015, Nadolny and Gaff 2018). Here, we present evidence for an established population of *A. maculatum* in New York City and discuss its possible establishment in New Jersey.

## Materials and Methods

### New York

The New York City Health Department conducts an annual city-wide blacklegged tick surveillance program in city parks, coastal-marshes, and grasslands throughout five boroughs (Bronx, Brooklyn, Manhattan, Queens, and Staten Island) of the city (NYC Health 2016). In Staten Island, the only constituency of New York City where local transmission of tick-borne diseases is known to occur (DOHMH 2021), surveillance is performed in several local parks. Specifically, since 2018 multiple locations are routinely sampled within Freshkills Park **(Figure 2)**, a large grassland that was once the world’s largest landfill situated in the central-west region of the island, and which now comprises a diverse habitat of 2,200 acres of grass and brushland (NYCDPR 2019). Freshkills Park is on the Atlantic flyway and several new bird species have recently established in the area and replaced the seagulls (Laridae), which in the past were the most common birds (TFPA 2016).

**Figure 2.**
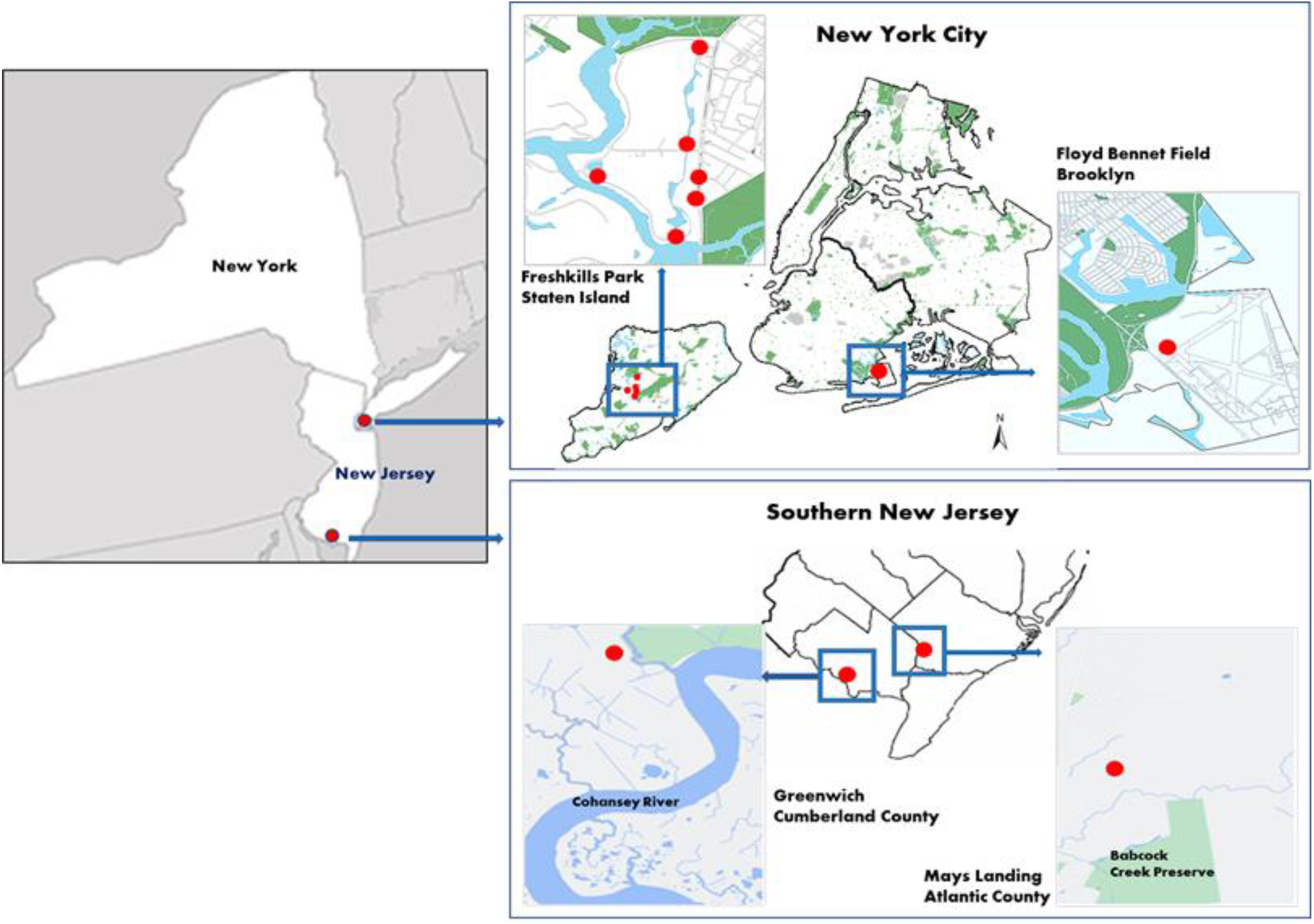
Surveillance for *A. maculatum* in New York City and New Jersey. The red dots in the two detailed maps on the right are sites where *specimens of A. americanum* were collected.

All surveys were performed using a tick drag made of a 1-m^2^ piece of white corduroy cloth pulled behind the operator over the ground and through low vegetation for 200 meters (Ferrell and Brinkerhoff 2018). The drag cloth was inspected every 20 m and attached ticks were removed with forceps and stored in 95% ethanol in labeled 1.5 ml plastic tubes. In the laboratory the ticks were identified to species using morphological keys (Pratt and Littig 1962, Keirans and Durden 1998) and a subset of specimens were confirmed using DNA barcoding (see below).

### New Jersey

Except for Monmouth Co., where a passive tick surveillance program was initiated in 2006 and active surveillance in 2017 (Jordan and Egizi 2019) routine tick surveillance is not performed in NJ. However, NJ has a strong interagency partnership between local and state entities involved in vector and vector-borne disease surveillance that facilitated the efforts described below.

On August 31, 2018, a resident of Mays Landing, Atlantic County NJ removed a tick from their dog (*Canis familiaris*) and submitted a photograph to the University of Rhode Island’s Tick Spotters website (https://web.uri.edu/tickencounter/tickspotters/submit/), who advised the resident to contact the Rutgers Center for Vector Biology (CVB, https://vectorbio.rutgers.edu). The resident forwarded the photograph of the tick by email to D.M.F. and eventually shipped the specimen therefore allowing formal identification. Using both morphology (Keirans and Litwak 1989) the specimen was identified as an adult male *Amblyomma maculatum*. The photograph elicited a series of immediate tick surveys of neighboring areas (gas pipeline right-of-way on Rt. 50) and parks (Atlantic County Military Park and Clover Leaf Park on Pinehurst Drive) on September 1-2, 2018. Although September is the tail end of adult *A. maculatum* activity (Maestas et al. 2020) additional surveillance was conducted on September 12, 2018, including the neighborhood around the resident’s home and coastal sites near the Egg Island Wildlife Management Area. Sites were chosen based on apparent suitability of habitat for *A. maculatum* obtained from a literature review, advice from experts, and accessibility. Surveys were developed using tick sweeps made of white 1-inch PVC pipe and a 0.5 m x 0.5 m white crib flannel (Egizi et al. 2019). Adult ticks were identified by experienced collectors in the field, but nymphs and larvae were collected with painter’s tape and placed in a labeled plastic bag for later examination in the laboratory. Ticks preserved in 80% ethanol were also obtained from nearby veterinarian practices on September 11, 2018.

Follow up surveillance efforts continued in 2019: In June 17-19, 2019, coastal areas of Cape May County, NJ were surveyed for *A. maculatum* focusing on coastal grassland habitats like those described in Maestas et al. (2020). On August 12, 2019, surveys were conducted in Cumberland Co. NJ, beginning where Tindall Island Rd. abuts the salt marsh (39.367496, −75,362600) and extending to grasslands and ecotones near the mouth of the Cohansey River, Dix Wildlife Management Area and Nantuxent Wildlife Management Area as these areas were directly across the bay from established populations of *A. maculatum* in Delaware (Florin et al. 2014).

### Barcode identification

A fragment of cytochrome oxidase 1 (*cox1*, the barcoding locus) was sequenced from a subsample of specimens identified as putative *A. maculatum* based on morphology. For adults, a single leg was removed and extracted using either a Qiagen DNeasy Blood & Tissue kit (Qiagen Inc., Valencia, CA) or a modified HotSHOT protocol (Truett et al. 2000, Egizi, Bulaga-Seraphin, et al. 2020). For larvae, DNA was also extracted using a Qiagen DNA extraction kit, after homogenizing the larval tissues with a 5mm stainless steel bead in Buffer ATL. Genomic DNA was eluted in 50ul Buffer AE. The barcoding locus *cox1* was amplified with Qiagen HotStarTaq Master Mix (Qiagen Inc., Valencia, CA) and primers HCO2198/LCO1490 (Folmer et al. 1994). The 25 μl reaction volume contained 12.5 μl HotStarTaq 2 x Master Mix, 1ul each primer (10μM), 1μl template DNA, and the remainder ultrapure water. PCR products were visualized on a 2% agarose electrophoresis gel, then primers and unincorporated dinucleotides were removed with ExoSAP-IT (USB Corporation, Cleveland, OH) and the DNA fragments were Sanger sequenced in both directions (Genewiz, South Plainfield, NJ). Sequences were assembled and trimmed in Geneious 10.2.3 (Kearse et al. 2012) and compared with known tick *cox1* sequences in a NCBI BLAST search (https://blast.ncbi.nlm.nih.gov/Blast.cgi).

### Testing of adults for *Rickettsia parkeri*

Ten adult *A. maculatum* ticks collected from Staten Island (NYC) were tested for the presence of *R. parkeri* DNA by PCR. Two types of qualitative PCR assays were employed for detection of *R. parkeri*. For genus-level detection, we used a primer pair that was designed to detect a fragment within the *ompB* gene of the most common spotted fever group *Rickettsia* species (Sanchez-Vicente et al. 2019) (Table 1).

**Table 1.**
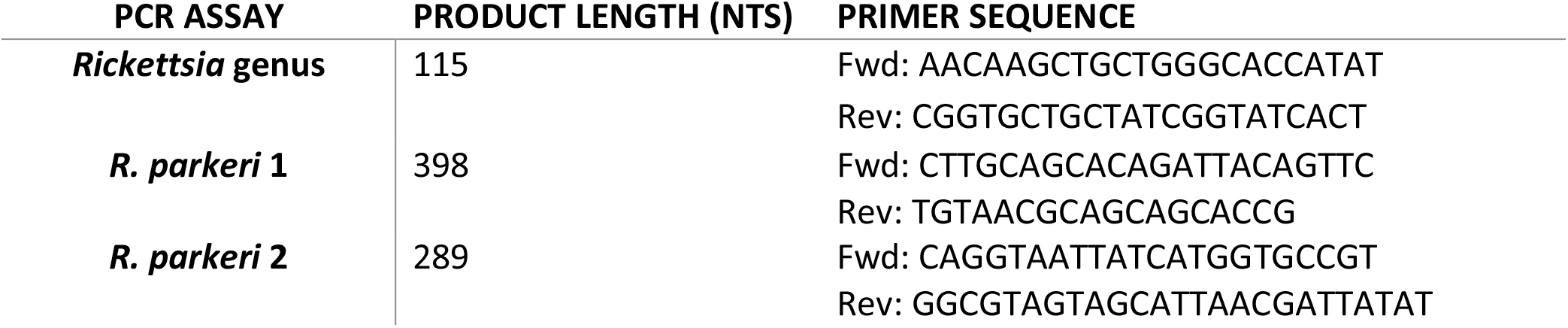
PCR assays and primers used for detection of *Rickettsia parkeri* in *A. maculatum* adults.

For specific *R. parkeri* detection, *ompB* sequences from multiple *Rickettsia* species available in GeneBank were aligned using Geneious 10.0.9 software. This included two *R. parkeri* sequences (accession numbers AF123717 and KF660535) as well as sequences from *R. buchneri* (EF433951), Candidatus *R. andeanae* (GU395297), *R. monacensis* (JX625150 and LN794217), *Rickettsia* sp. Bar29 (AF123710), *R. hulinensis* (AY260452), *R. massiliae* (CP000683), *R. amblyommatis* (CP015012), *R. raoultii* (CP019435), *R. japonica* (CP032049), *R. aeschlimannii* (KU961544), *R. rickettsii* (GU395293), *R. felis* (AF182279), *R. helvetica* (MF163037), *R. akari* (AF123707), *R. typhi* (L04661), and *R. prowazekii* (AF161079). Two primer sets (*R. parkeri* 1 and *R. parkeri* 2 – Table 1) were selected based on in silico specificity for *R. parkeri* and the desired length of the PCR product (>250 nt).

Prior to nucleic acid extraction, each individual *A. maculatum* was washed twice with 1 ml nuclease free water and homogenized using a 21-gauge, 1.5-in needle in 50 μl of nuclease free water. The homogenate was added to NucliSENS® easyMAG® lysis buffer and total nucleic acids were extracted according to manufacturer’s protocol on the easyMAG platform (bioMérieux, France) and eluted in 40 μl of elution buffer.

All PCR assays were performed in a 25 μl-reaction mixture with 2 μl of template, 0.2 μM of the forward and reverse primer, 1 μl of GC enchancer and 12.5 μl of AmpliTaq Gold 360 Master Mix (Applied Biosystems). PCR reaction conditions for each assay consisted of: 95°C for 10 min, 40 cycles of 95°C for 30 s, 55°C for 45 s, and 72°C for 30 s, followed by 10 mins at 72°C. PCR products were resolved on a 2% agarose gel in Tris-borate-EDTA (TBE) buffer. PCR products were sequenced by Sanger sequencing, analyzed using Geneious 10.0.9 software, and identified as *R. parkeri* through BlastN homology searches.

### Testing of larval ticks for *Rickettsia* spp

Twenty-four larvae collected in August 2020 were tested for *Rickettsia* spp. using a species-specific qPCR duplex targeting *R. parkeri* and *R. rickettsii* as described in Egizi et al. (2020) with primers and probes adapted from published assays (Jiang et al. 2005, 2012, Gaines et al. 2014). They were also tested with a broader spotted fever group qPCR assay for the 17-kDa gene (Jiang et al. 2012) as previously described (Egizi, Gable, et al. 2020). To confirm positive samples, conventional PCR assays for spotted fever rickettsia were used to sequence fragments of the outer membrane protein A (*ompA)* gene with primers Rr190.70p and Rr190.701 (Regnery et al. 1991, Roux et al. 1996) and the citrate synthase (*gltA)* gene with primers RrCS372 and RrCS989 (Kollars and Kengluecha 2001). PCRs were performed in a 25ul reaction volume with 12.5 ul Qiagen HotStarTaq Master Mix (Qiagen Inc., Valencia, CA), 1ul each primer (10um), 2ul template DNA, and the remainder ultrapure water. PCR products were visualized on a 2% agarose electrophoresis gel, then primers and unincorporated dinucleotides were removed with ExoSAP-IT (USB Corporation, Cleveland, OH) and the DNA fragments were Sanger sequenced in both directions (Genewiz, South Plainfield, NJ). Sequences were assembled and trimmed in Geneious 10.2.3 (Kearse et al. 2012) and compared with known *Rickettsia* sequences in a NCBI BLAST search (https://blast.ncbi.nlm.nih.gov/Blast.cgi). For all reactions, positive controls used were two adult *A. maculatum* also from Staten Island that were extracted for DNA barcoding but found by conventional PCR assays to be infected with *R. parkeri* (100% identity with Accession #U43802 for *ompA* and #U59732 for *gltA*).

## Results

### New York

The first *A. maculatum* from New York City was a male collected on July 18, 2018 from Freshkills Park, Staten Island at the geographic coordinates (X,Y): 40.574051, −74.170539. In 2019, we obtained another male and a female at a different location in the same park (40.573836, −74.181022). The male (accession# USNMENT986148) was crawling on a field staff member on July 19 and we collected the female (accession# USNMENT986147) on July 25 by dragging at the same location. In 2020, we collected the first adult *A. maculatum* (a female) on June 22 (40.57193, −74.170673), and when we surveyed for ticks on July 16 at four different locations (40.5677, −74.1727; 40.57766, −74.1715353; 40.58821, −74.170052; 40.57193; −74.170673) within Freshkills Park we collected 15 adult *A. maculatum* (10 females and 5 males). Due to stringent COVID pandemic restrictions in New York City at the time, we were only able to sample one location (40.57193; −74.170673) on August 27 when we collected 2 females and 66 larvae of *A. maculatum*. One of the larvae has accession# USNMENT986149. In summary, in 2020 we collected a total of 18 adults and 66 larvae from Freshkills Park.

The dominant plant species at Freshkills Park (**Figure 3A)** are switchgrass (*Panicum virgatum* L.), Indian nut grass (*Sorghastrum nutans* (L.) Nash), roundfruit rush (*Juncus compressus* Jacq.), and big bluestem (*Andropogon gerardii* Vitman) (TFPA 2019). Some less common plant species found in Freshkills Park are blue grama (*Bouteloua gracillis*), gama grass (*Tripsacum dactyloides*), blue-green grass (*Bouteloua curtipendula)*, little bluestem (*Andropogon scoparium*), purple lovegrass (*Eragrostis spectabilis*) and several flowering weeds including blacked-eyed susan (*Rudbeckia hirta*), and butterfly weed (*Asclepias tuberosa*) (TFPA 2015).

**Figure 3.**
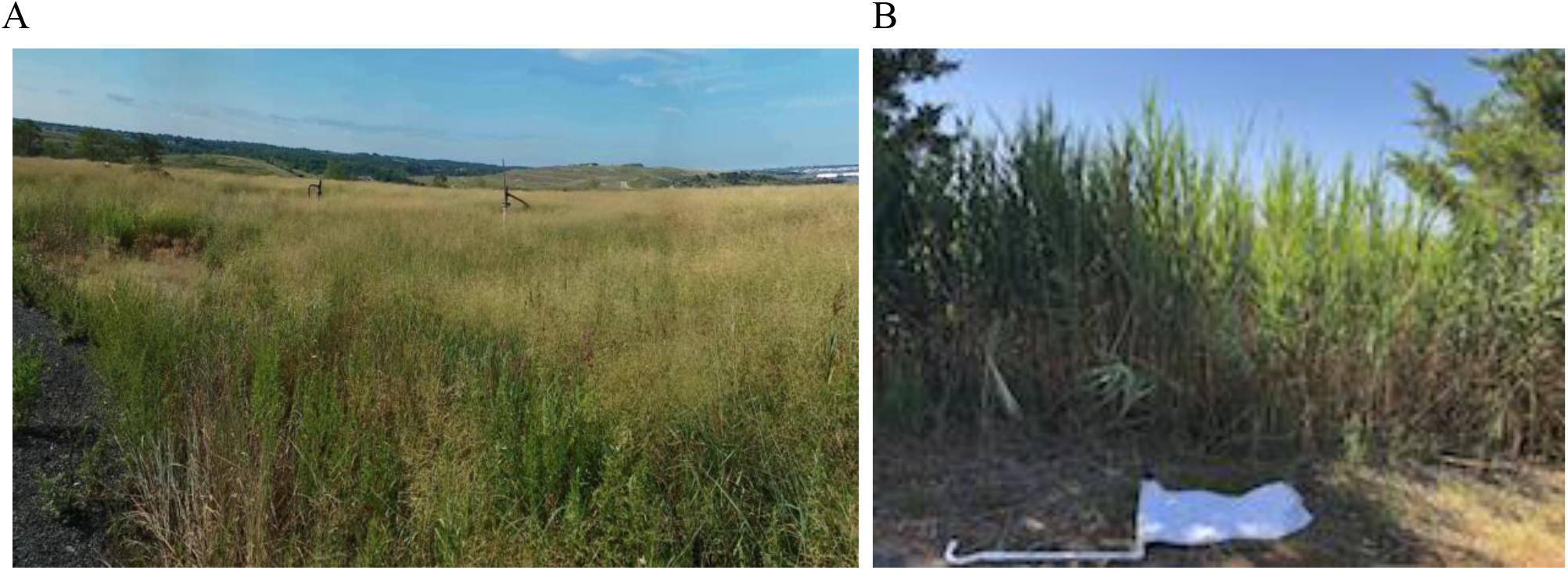
**A**. Representative vegetation and landscape configuration at Freshkills Park, Staten Island, NY. **B**. Invasive *Phragmites* in Tindall Island, Cumberland Co. NJ from where the female *A. maculatum* was collected; the tick sweep used for sampling is included for scale.

DNA barcoding was used to confirm the identification of one adult female and one adult male collected in Staten Island on July 16, 2020, and eight larvae collected on August 27, 2020. The male and female had the same COI haplotype and matched NCBI Accession #KU302500 (*Amblyomma maculatum* from Georgia, USA (Lado et al. 2018)) with 100% identity. All eight larvae had the same haplotype as each other, but 1 bp distinct from the adults, matching several NCBI Accessions (#MG251316, MG251314, MG251313, KU302504, KU302503, KU302499, KU302497, and KU302495 from Arizona, Georgia, and Florida, USA (Lado et al. 2018)) with 100% identity.

Outside Staten Island, we collected two female *A. maculatum* on July 14, 2020, from Floyd Bennet Field (40.59405, −73.901726) in the borough of Brooklyn (New York City) during our regular drag sampling for blacklegged ticks. Floyd Bennett Field is in southeast Brooklyn along the shore of Jamaica Bay. Its natural area contains over 1,300 acres of grassland, salt marshes, and tidal mudflats. At Floyd Bennett Field, open areas are dominated by native species including grasses like little bluestem (*Schizachyrium scoparium* (Michx.) Nash) and phragmites (*Phragmites australis* (Cav.) Trin. ex Steud. (ITIS)) (NPS 2021).

### New Jersey

The male tick removed by a resident of Mays Landing, Atlantic Co., NJ (39.465161, −74.713194) from a dog was confirmed to be *Amblyomma maculatum*n by DNA barcoding (closest match to KX360344 and KM839245 from Ontario, Canada, with 99.8% identity). We used a single leg for the barcode and submitted the remainder of the specimen as a voucher to the US National Tick Collection (accession# USNMENT1512539). Because its COI sequence differed by 1 bp from publicly available sequence we have submitted it to GenBank (Accession # pending). Conversations with the resident determined that neither the resident nor her dog had traveled outside NJ for several weeks before the tick was found. Subsequent investigations of the resident’s neighborhood, nearby parks, as well as ticks collected from veterinary clinics in the area, detected established populations of *A. americanum, Dermacentor variabilis* Say, and *I. scapularis*, but did not recover any additional *A. maculatum*. On August 12, 2019, we collected a female *A. maculatum* (USNMENT1512538) on Tindall Island, Cumberland Co. NJ (39.367496, −75,362600) off invasive *Phragmites* on the side of Tindall Island Rd (**Figure 3B**). This was the only specimen of *A. maculatum* we detected in NJ that year and since.

### Pathogen testing

A total of 5 out of 10 adult *A. maculatum* ticks tested positive for *R. parkeri* (50%) by all three *ompB* PCR screening assays and subsequent sequencing of PCR products. The sequences were identical from all five positive specimens, with 100% homology to *Rickettsia parkeri* str. Portsmouth (accession #: CP003341.1) that was isolated in cell culture from an eschar biopsy specimen and *Rickettsia parkeri* str. Maculatum 20 (accession # AF123717.1) isolated from ticks in Mississippi, USA.

Although none of the larval ticks were positive for *R. parkeri*, one larva out of the 24 tested (4.17%) was positive for *Rickettsia felis*. Sequencing of the PCR products revealed 100% identity with *ompA* (Accession #AF210694) and *gltA* (#AF210692) sequences of the *R. felis* type strain, isolate Marseille-URRWFXCal2(T) as well as sequences from North, Central and South America, Europe, Asia, and Africa (Raoult et al. 2001, La Scola et al. 2002).

## Discussion

The multiyear, multistage collections of the Gulf Coast tick, *Amblyomma maculatum* in Staten Island, New York City, their detection in Brooklyn, NYC as well as in Cape May Co., NJ and Cumberland Co. NJ, USA, indicate further northward spread of this species along the Atlantic coast. Of note, a specimen of *A. maculatum* was collected in 1981 from Cattaraugus County, NY (Wiedl 1981), which is near Niagara Falls, 260 miles northwest of the current 2018-2020 collection site but to our knowledge the species has not since been detected there. Detection of >6 individuals of a single life stage or > 1 life stage collected in a county within a 12-month period signifies an established tick population according to CDC guidance (https://www.cdc.gov/ticks/pdfs/Tick_surveillance-P.pdf). Applying these criteria to our data, *A. maculatum* would be considered established on Staten Island (Richmond County) although not in Brooklyn (Kings County) or in NJ. Additionally, there were 2 unique COI haplotypes among the barcoded *A. maculatum* samples, suggesting that the Staten Island population are not descendants of a single lineage and providing further evidence for an established (as opposed to adventive) population.

Biotic and abiotic features that are key determinants of the survival and establishment of *A. maculatum* in an area such as large grasslands in mid-to-late successional stages (Cilek and Olson 2000, Teel et al. 2010, Paddock and Goddard 2015, Nadolny and Gaff 2018) occur on Staten Island especially in Freshkills Park. Furthermore, flocks of migratory birds, traveling along the Atlantic flyway, rest for days at numerous stopover sites on the island during their spring migration(https://migbirdapps.fws.gov/) providing a means of ongoing tick introductions. Staten Island has abundant populations of resident hosts, including white-tailed deer, which is considered the main host of adult *A. maculatum*, and small mammals and birds to feed nymphs and larvae (Sonenshine 2018). In addition, daily maximum and minimum temperatures in NYC have risen 1.26°F and 1.98°F, respectively, since 1970 (NOAA 2021) making the present climate more hospitable to this southern species. Of note, several nonnative mosquitoes have recently established in the city (Bajwa 2018).

In NYC, NJ as well as in CT (Molaei et al. 2021), the detection of a crawling or feeding *A. maculatum* incentivized local entomologists to develop enhanced surveillance and in NYC and CT, but not in NJ, eventually detect established populations. Of note, the fact that in all states (including NJ) the first specimen of *A. maculatum* detected was a male is likely because male *A. maculatum* remain on the host much longer than females since they mate on host and try to mate with multiple females (Hooker et al. 1912). The detection of established populations of *A. maculatum* in NYC may be less surprising than their apparent absence from New Jersey (which lies directly between areas with established populations in NYC and Delaware). *A. maculatum* have also been sporadically collected from NJ’s western neighbor Pennsylvania although to date there are no known established populations (Pak et al. 2019).

The presence of established populations in coastal CT and now also in Staten Island suggests that the species should be able to thrive in NJ. The detection of two adult *A. maculatum* in NJ indicates the species is brought into the state. Therefore, our failure to detect established populations of *A. maculatum* in NJ may be more likely the result of inherent difficulties in detecting low density, spatially heterogeneous incipient populations (Mehta et al. 2007, Berec et al. 2015) compounded by the lack of standardized surveys for ticks across NJ (Egizi et al. 2019). As warming temperatures and milder winters may enable *A. maculatum* populations to grow and spread further into the northeast region, it is likely that established populations of *A. maculatum* will be detected in NJ in future years.

The presence of *A. maculatum* in NYC, CT and possibly in NJ is of significant public health importance. We detected *Rickettsia parkeri*, a tick-borne human pathogen, in 50% of adult *A. maculatum* ticks collected from Freshkills Park, Staten Island (NYC). These results concur with findings of other studies along the US east coast where prevalence of *R. parkeri* in field collected *A. maculatum* ticks can be very high, ranging from 20% to 55.7% (Fornadel et al. 2011, Varela-Stokes et al. 2011, Wright et al. 2011, Nadolny et al. 2014, Maestas et al. 2020, Molaei et al. 2021). They are, however, somewhat higher than the prevalence recorded from the two states closest to NYC, which recently detected *A. maculatum* populations: Delaware (21%) (Maestas et al. 2020) and Connecticut (29.6%) (Molaei et al. 2021). NYC and its surrounding areas (formally known as the New York-Newark Combined Statistical Area by the US Office of Management and Budget and comprising parts of NY, NJ, CT, and PA) is the most populous region in the US with 23.5 million residents as of the 2020 census (U.S. Census Bureau 2021). Staten Island alone contains 474,893 people and Freshkills Park experiences 3,000 visitors per year (personal communication: Richard Simon, Director, NYC Parks Department), a number that will be higher when all sections of the park are open to the public (Klenosky et al 2017) and all of which may now be at risk of contracting *Rickettsia* pathogens from *A. maculatum* tick bites.

Interestingly, *R. parkeri* was not detected in the larvae tested despite high rates of adult infection and known transovarial transmission. However, as the larvae were all collected in the same area on the same day (and all 8 barcoded larvae shared the same COI haplotype) it is likely that they were all from the same egg mass from an uninfected female. Instead, one larva was infected with *Rickettsia felis*, a pathogen whose primary vector and reservoir is the cat flea (*C. felis*) although it has been detected in a range of other arthropods, including other species of fleas, mosquitoes, and ticks (Brown and Macaluso 2016). In US ticks previously *R. felis* was detected in *A. maculatum* from Virginia and Mississippi (Jiang et al. 2012) and a questing *H. longicornis* in Virginia (Thompson et al. 2021) however the involvement of these species in transmission of *R. felis* is unknown. To our knowledge *R. felis* has not been reported from arthropods or humans in NYC, although Hoque et al. (2020) detected the bacterium in a blood sample from a domestic cat in Brooklyn.

The arrival of *A. maculatum* brings a novel risk of tickborne illnesses to the NY metropolitan area, a risk only expected to increase as the species further expands its range and becomes more abundant. Physicians in the area should be informed of the possibility of human infection with *R. parkeri* and *R. felis* associated illnesses and veterinarians should be apprised of possible canine hepatozoonosis. Visitors to Freshkills and other parks in the region should be reminded to take precautions against tick bites. Lastly, neighboring areas like NJ should continue to monitor for the establishment of this species.

## Acknowledgements

We would like to acknowledge James Occi for jumping into action immediately after the first *A. maculatum* was detected in NJ and developing extensive surveillance in Mays Landing parks and later, with Alvaro Toledo, wildlife areas in coastal Cape May. We are also indebted to Kim Cervantes from the NJ Department of Health and John (Doug) Abdill from the Atlantic County Mosquito Control Program for contacting local veterinarians and obtaining ticks collected from dogs in Mays Landing in August 2018. We thank Alvaro Toledo, Francisco Ferreira, Andre Bhandoola and Raoul Bhandoola for trying unsuccessfully to collect *A. maculatum* in June 2019 across southern NJ. They kept the spirits high even while experiencing an assault of lone star tick nymphs. The NJ work was partly supported by State of New Jersey FY22 Tick Research and Control - Special Purpose Funding to NJAES/CVB.

